# Predicting intensities of Zika infection and microcephaly using transmission intensities of other arboviruses

**DOI:** 10.1101/041095

**Authors:** Isabel Rodríguez-Barraquer, Henrik Salje, Justin Lessler, Derek A.T. Cummings

**Author notes:** Correspondence: Isabel Rodríguez-Barraquer.

## Abstract

The World Health Organization has declared Zika Virus a Public Health Emergency of International Concern due to the virus’ emergence in multiple countries globally and the possible association of Zika virus with microcephaly and neurological disorders. There is a clear need to identify risk factors associated with Zika infection and microcephaly in order to target surveillance, testing and intervention efforts. Here, we show that there is a strong correlation between the incidence of Zika in Colombian departments and the force of infection (but not the crude incidence) of dengue, a virus transmitted by the same mosquito species, *Aedes aegypti* (R^2^ = 0.41, p<0.001). Furthermore, we show that there is also a strong correlation between the incidence of microcephaly in Brazilian states and the force of infection of dengue (R^2^ = 0.36, p<0.001). Because dengue, Zika and chikungunya are transmitted by the same vector, these associations provide further support to the supposition that Zika virus infection during pregnancy causes microcephaly. In addition, they provide an opportunity to project the expected incidence of microcephaly in multiple dengue endemic locations across Colombia and the American continent. Detailed knowledge of dengue transmission should be use to target efforts against Zika and other flaviviruses.

## Introduction

The World Health Organization has declared Zika virus a Public Health Emergency of International Concern due to the virus’ emergence in multiple countries globally and the possible association of Zika virus with microcephaly and neurological disorders.^1-3^ As of February 2016, Brazil has had the overwhelming majority of microcephaly associated with Zika virus though the causal link between Zika virus and microcephaly remains unproven^4,5^. Colombia has reported significant numbers of Zika virus infections, but has not yet reported a significant increase in microcephaly cases.^6^ There is a clear need to identify risk factors associated with Zika infection and microcephaly in order to target surveillance, testing and intervention efforts.

Dengue virus, transmitted by the same species of mosquito as Zika, has caused considerable disease burden in both countries over the past two decades.^7-9^ More recently chikungunya, another arbovirus transmitted by the same vector, was introduced into the continent and caused large outbreaks.^10,11^ As these three pathogens share a vector, it is expected that metrics of the transmission potential of dengue and chikungunya should be correlated with the incidence of Zika virus if mosquito density, climate and other environmental factors affect transmission of all three pathogens similarly. Moreover, if Zika virus infection during early stages of pregnancy is associated with neurological malformations such as microcephaly, the transmission intensity of other *Aedes* transmitted viruses (including dengue and/or chikungunya) can potentially be used to assess a region’s microcephaly risk.

Here, we model the correlation between transmission intensity of dengue and chikungunya in Brazilian states with reported microcephaly incidence. We also correlate dengue and chikungunya transmission intensity in Colombia to Zika incidence, as no microcephaly cases have been reported up to now. Furthermore, we build on the observed correlation to map the expected incidence of microcephaly in multiple dengue endemic locations across Colombia and the American continent that have reported autochthonous Zika transmission but no microcephaly cases yet.

## Methods

We obtained data on the number of reported cases of microcephaly in each state of Brazil from the weekly reports released by the Brazilian health ministry.^12^ Since no cases of microcephaly have been reported in Colombia as of now, we obtained data on the number of Zika cases reported by each department. ^6^ In addition, we obtained data on the number of dengue and chikungunya cases reported by Brazil (chikungunya 2014-2015; dengue 2002-2012) and Colombia (chikungunya 2014-2015, dengue 2007-2012), and census data from each country during the observed period ^13 14,15^

To assess whether metrics of transmission of the three arboviruses are associated, we correlated the incidences of microcephaly/Zika for each department in Brazil and each department in Colombia to the incidences of dengue (average over the 5 last years of data available) and chikungunya during the 2014-15 outbreak in those departments. Taking into account that dengue is highly endemic in both Brazil and Colombia and a large proportion of the population is no longer susceptible, we also correlated the incidence of microcephaly/Zika to the mean transmission hazard, i.e., the mean force of infection (FOI), of dengue. The force of infection is a metric of incidence among the susceptible population, and is a more robust metric of transmission potential. Estimates of the force of infection of dengue were derived for each state/department of Colombia and Brazil, by fitting catalytic models to the age-stratified dengue incidence data from as has been previously described. ^16,17^

All analyses were performed in R version 3.2. ^18^

## Results

We find that both the incidence of microcephaly in Brazil and the incidence of Zika in Colombia are strongly correlated with the force of infection of dengue (Figures 1, coefficient of determination R^2^ = 0.36 and 0.41, respectively, p<0.001), but not with the recent incidence of dengue (Figure 1, R^2^ = 6e-5, p=0.97 and R^2^ = 0.01, p=0.55, respectively) or chikungunya (Figure 2, R^2^ = 0.01, p=0.54 and R^2^ = 0.11, p=0.09, respectively).

**Figure 1:**
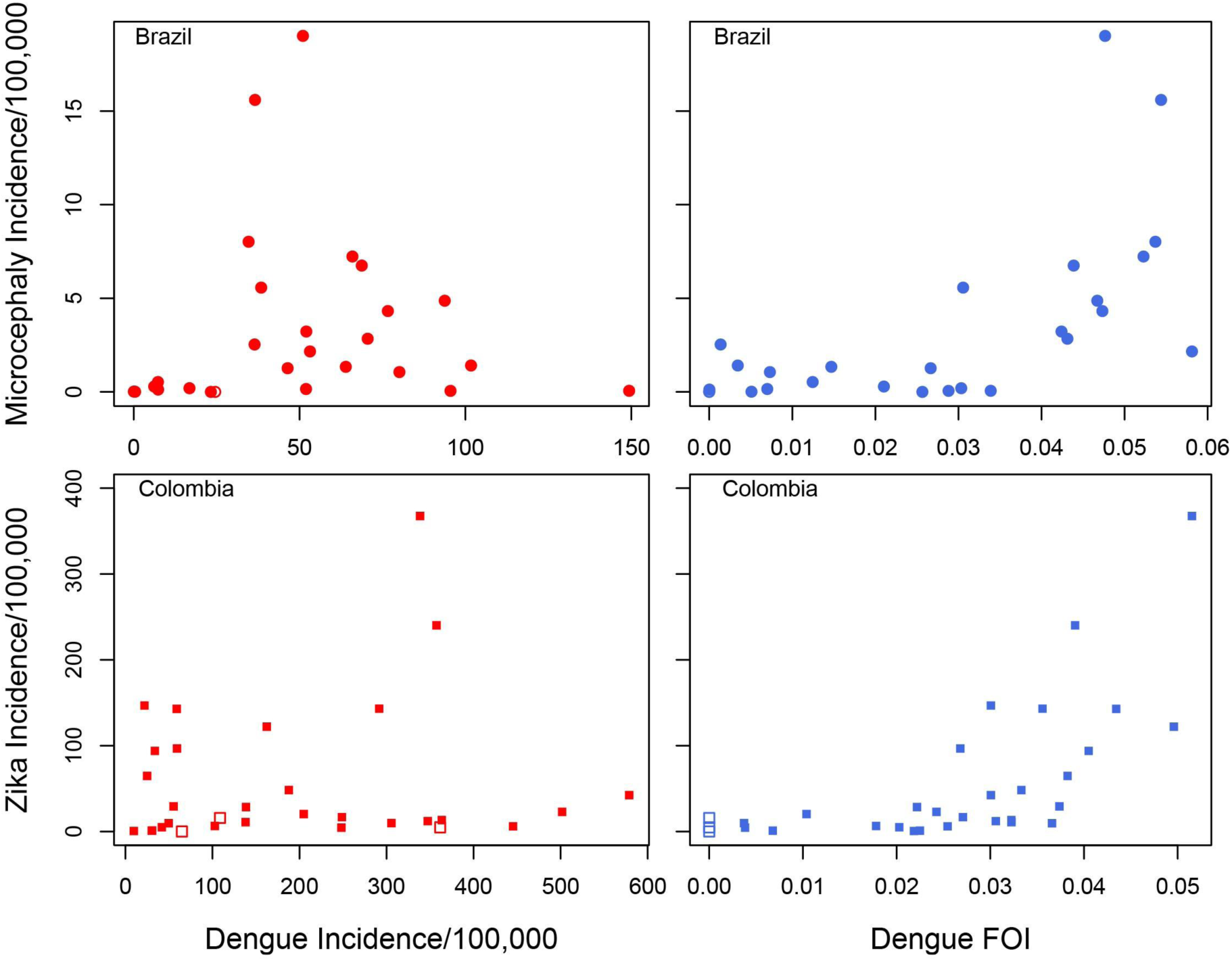
Correlation between dengue incidence (left) and dengue force of infection (right) with the incidence of microcephaly in states of Brazil and the incidence of Zika in continental departments Colombia. While there is no apparent correlation between the dengue incidence and either microcephaly or Zika (R^2^=6e-5and 0.01, respectively), a significant positive correlation is observed between the estimated force of infection and microcephaly/Zika incidence in the two countries (R2= 0.36 and 0.41, respectively. p<0.001). Open symbols indicate states/departments where low dengue cases counts didn’t allow estimation of the force of infection

**Figure 2:**
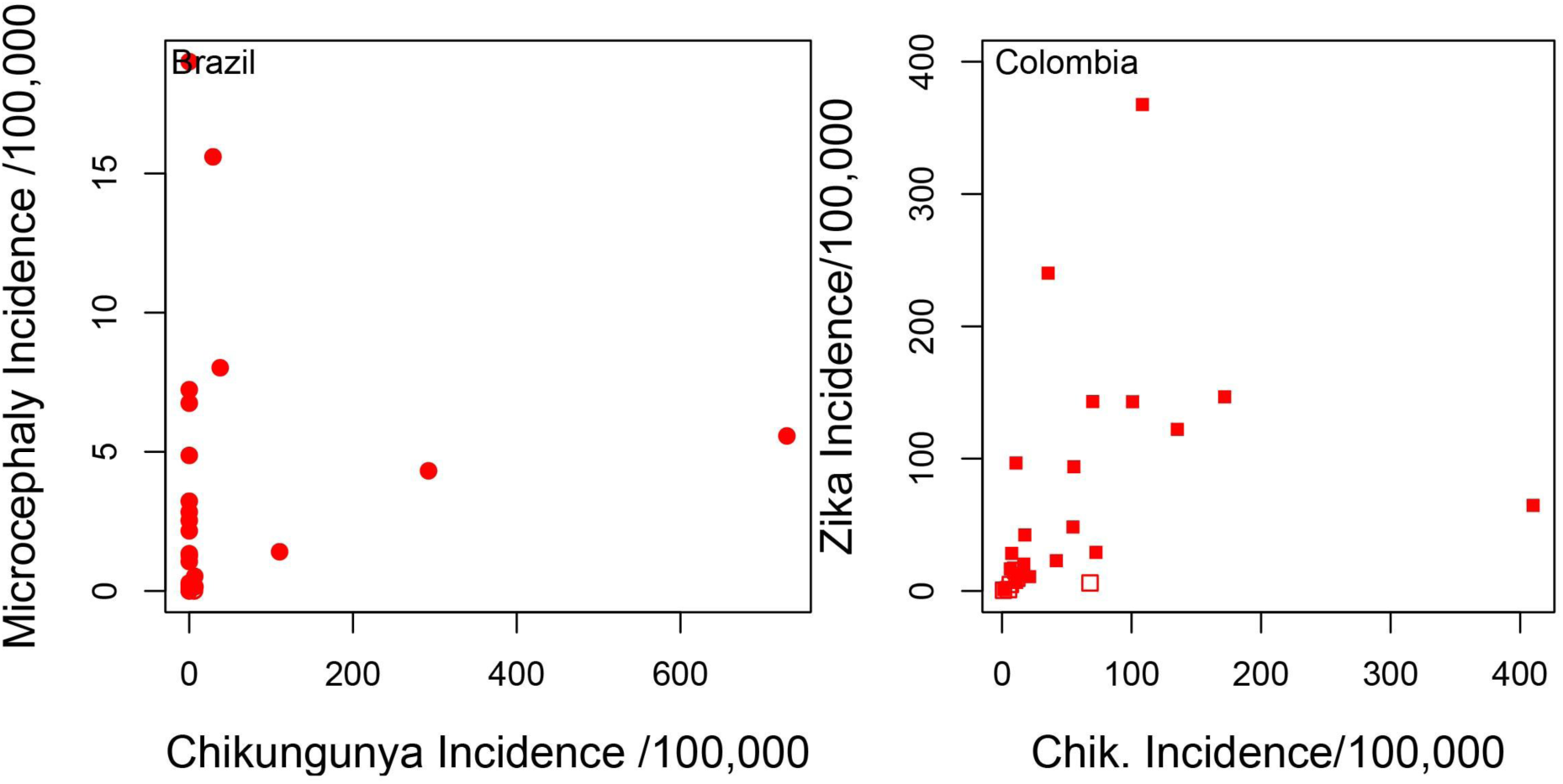
Correlation between Chikungunya incidence with the incidence of microcephaly in states of Brazil and the incidence of Zika in departments Colombia. There is only a weak correlation between Chikungunya and Zika incidence in Colombia (R^2^ = 0.11, p=0.09), and no observable correlation in Brazil (R^2^ = 0.01, p=0.54),.

**Figure 3:**
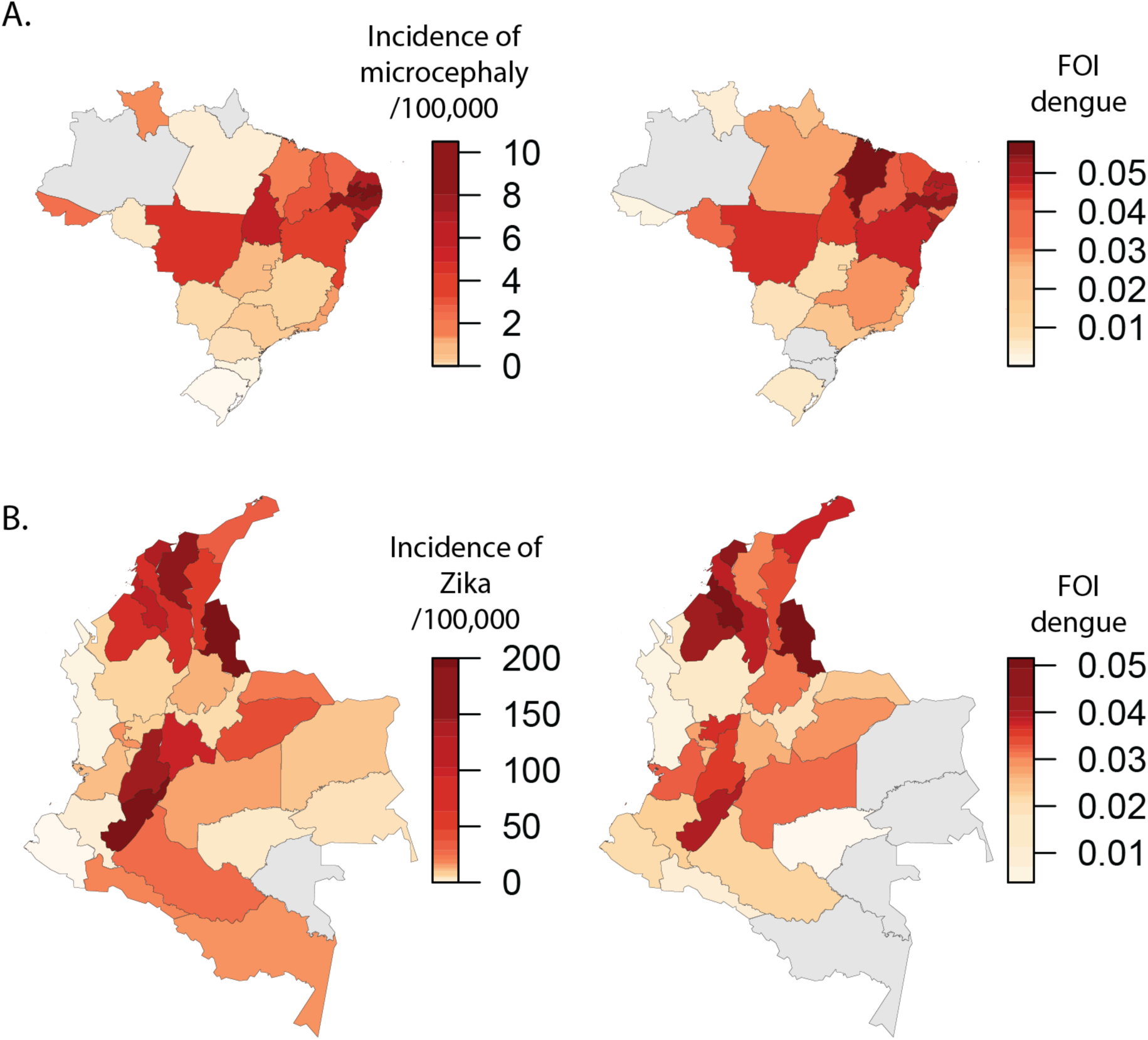
Maps comparing the incidence of microcephaly in states of Brazil (A) and the incidence of Zika in departments Colombia (B) to the estimated dengue force of infection.

With the exception of Paraiba and Pernambuco, that have reported incidences of microcephaly that are significantly higher than the rest of the country, there seems to be a linear association between dengue force of infection and incidence of microcephaly as of January 2016. Most of the microcephaly incidence seems to be concentrated in states where >3% of the susceptible population gets infected by dengue each year. In contrast, there is no apparent correlation between recent dengue incidence and microcephaly. Several of the states with highest incidence of microcephaly have reported relatively low dengue incidences over the past few years.

There is also an approximately linear association between the dengue force of infection and the incidence of Zika recorded in the continental departments of Colombia since it’s introduction in in October 2016. Each percent increase in the annual force of infection of dengue corresponds to a 44 per 100,000 (95%CI 24-64) increase in the incidence rate of Zika. While it was not possible to obtain reliable estimates of the dengue force of infection for 5/32 departments of Colombia, due to low case case-counts, the incidence of Zika reported in 4/5 of these states has been almost negligible. The island of San Andrés is the only exception to this trend, as it has reported the highest incidence of Zika in the country in this ongoing epidemic. Similar to Brazil, most of the incidence of Zika is concentrated in departments where the force of infection of dengue is 3% or higher per year.

To quantify the expected magnitude of the microcephaly epidemic in other settings that have not yet reported or observed cases, we mapped estimates of the dengue force of infection for multiple locations in Colombia to the observed association between dengue and microcephaly in Brazil (Figure 4, Table 1). For reference, we also mapped estimates of the force of infection of dengue in other Latin American settings derived from the literature^19^. While the absolute numbers can change as cases continue to be reported and investigated in Brazil, results from this mapping exercise suggest that, if the suspected causal link between Zika and microcephaly is real, the departments of Colombia that have experienced the highest incidences of Zika could expect incidences of microcephaly similar to those that have been observed in northeastern Brazil. However, given the difference in population size between the two countries, these incidences should result in a considerably lower number of microcephaly cases. If the relationship between dengue FOI and microcephaly incidence seen in Brazil holds in Colombia, we should expect to see 1024 cases (95%CI 440-1645) of Zika associated microcephaly to be reported over the first 5 months (of the microcephaly epidemic). Adjusting for the false positivity rates reported to date^20^ yields numbers that are more optimistic (387; 95CI% 166-621).

**Figure 4:**
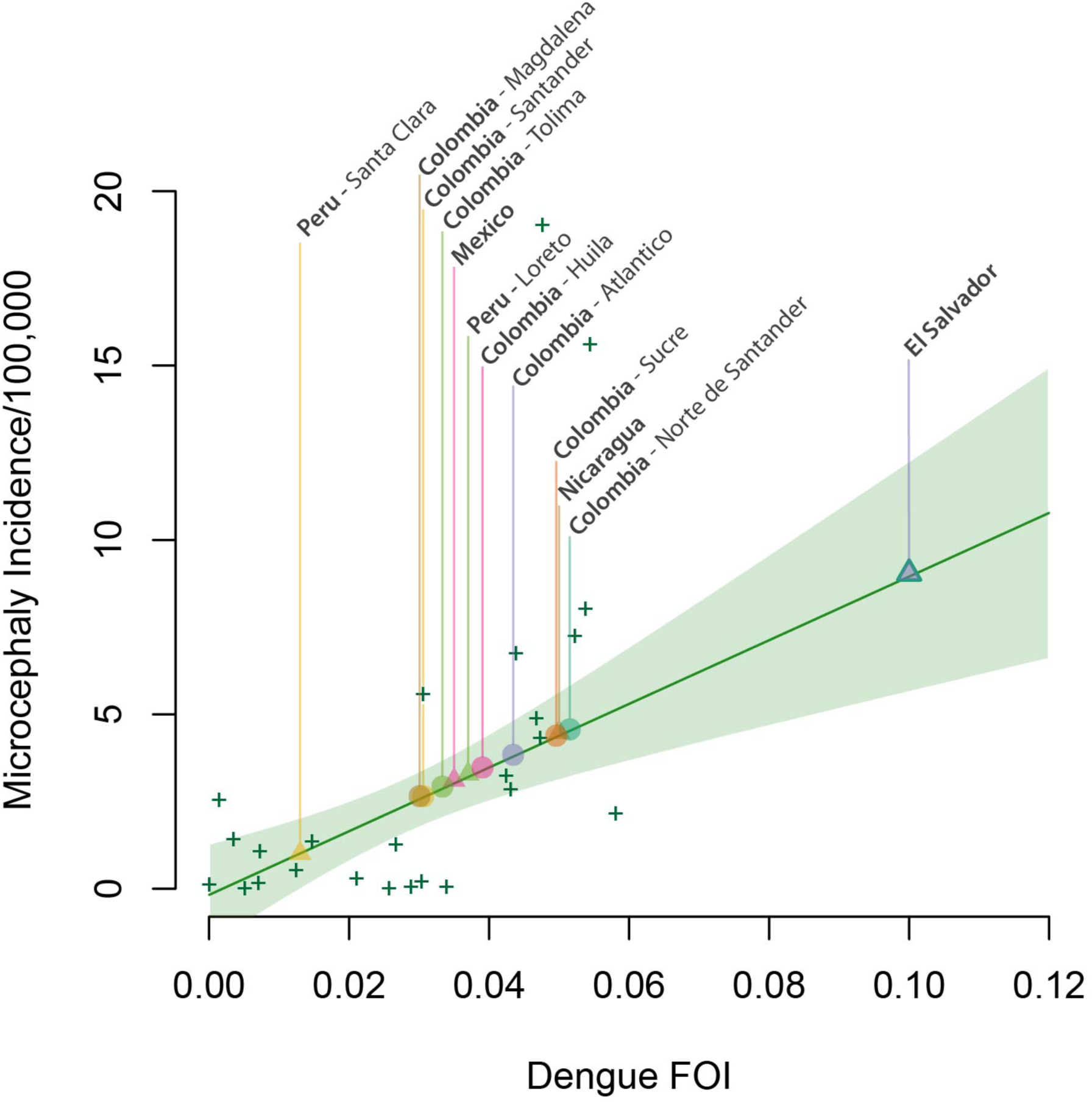
Mapping the expected microcephaly incidence (per 100,000 population) across different locations in Colombia, based on the dengue force of infection. The linear association (green line) between the dengue FOI and the incidence of microcephaly in Brazil was obtained by fitting a regression model to state level microcephaly/FOI data (shown as green “+”). The two outlying states with the highest microcephaly incidence (corresponding to Pernambuco and Paraiba) were excluded when fitting the model. Shaded area indicates 95% confidence bounds of this mean association. Symbols on top of the line map forces of infection estimated for multiple departments in Colombia (circles) to the expected microcephaly incidence. For reference, estimates of the force of infection for several locations in the Americas, derived from the literature are also shown (triangles). ^19^

**Table 1:**
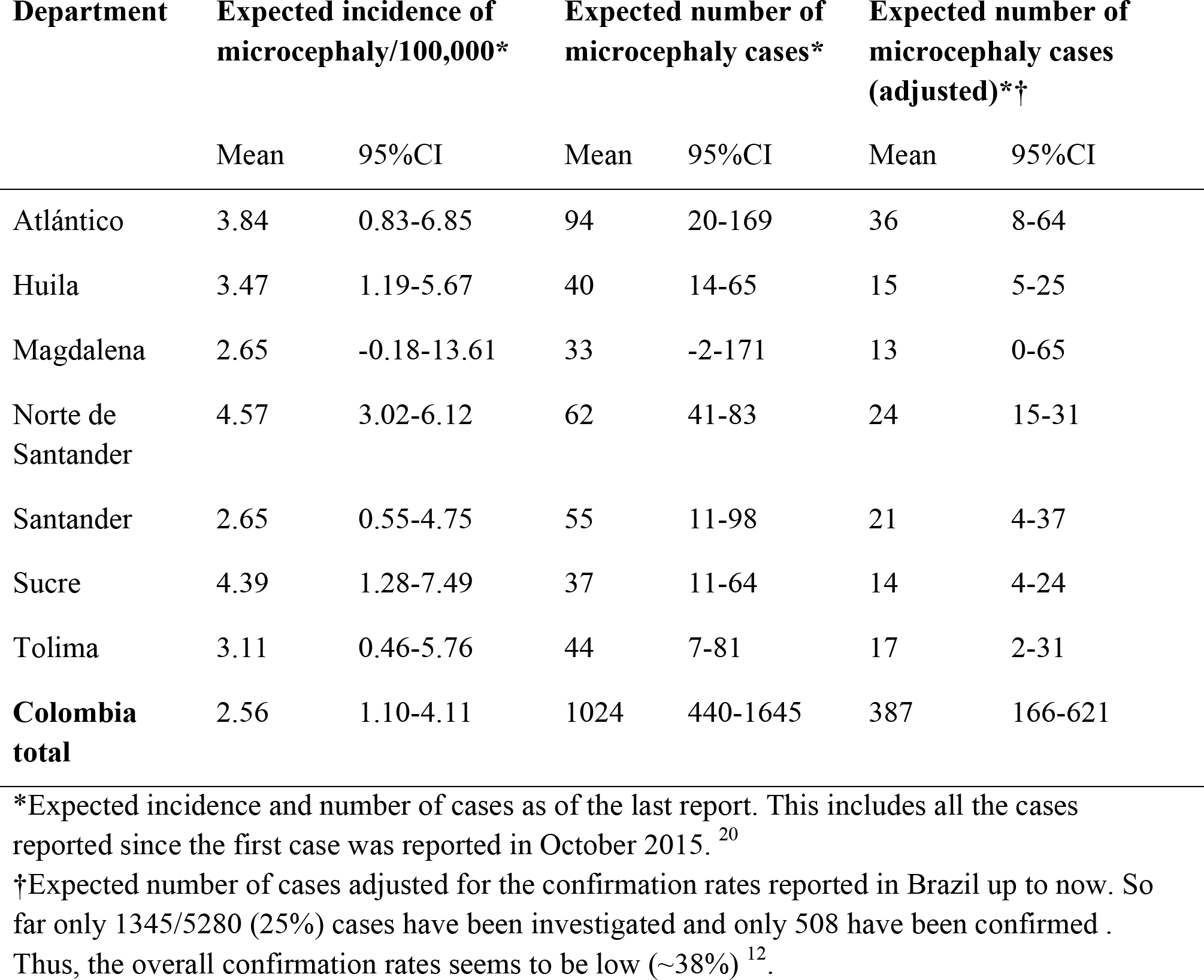
Expected incidence and number of microcephaly cases in selected departments of Colombia and in the country as a whole.

| Department | Expected incidence of microcephaly/100,000* |  | Expected number of microcephaly cases* |  | Expected number of microcephaly cases (adjusted)*† |  |
| --- | --- | --- | --- | --- | --- | --- |
|  | Mean | 95%CI | Mean | 95%CI | Mean | 95%CI |
| Atlántico | 3.84 | 0.83-6.85 | 94 | 20-169 | 36 | 8-64 |
| Huila | 3.47 | 1.19-5.67 | 40 | 14-65 | 15 | 5-25 |
| Magdalena | 2.65 | -0.18-13.61 | 33 | -2-171 | 13 | 0-65 |
| Norte de Santander | 4.57 | 3.02-6.12 | 62 | 41-83 | 24 | 15-31 |
| Santander | 2.65 | 0.55-4.75 | 55 | 11-98 | 21 | 4-37 |
| Sucre | 4.39 | 1.28-7.49 | 37 | 11-64 | 14 | 4-24 |
| Tolima | 3.11 | 0.46-5.76 | 44 | 7-81 | 17 | 2-31 |
| **Colombia total** | 2.56 | 1.10-4.11 | 1024 | 440-1645 | 387 | 166-621 |

\*Expected incidence and number of cases as of the last report. This includes all the cases reported since the first case was reported in October 2015. <sup>20</sup>
†Expected number of cases adjusted for the confirmation rates reported in Brazil up to now. So far only 1345/5280 (25%) cases have been investigated and only 508 have been confirmed . Thus, the overall confirmation rates seems to be low (~38%) <sup>12</sup>.

## Discussion

Here, we show that the incidence of Zika in Colombian departments and the incidence of microcephaly in Brazilian states are strongly correlated with the hazard of dengue, another arbovirus transmitted by mosquitos of the genus *Aedes*. These findings constitute further circumstantial evidence for the causal link between Zika infection and microcephaly incidence and provide a simple method that countries can use to assess their risk.

The correlation between Zika incidence (Colombia) and dengue forces of infection could be attributable to several mechanisms. The simplest is that areas with high suitability for transmission of dengue are also highly suitable for Zika virus transmission, mediated through multiple vector, human host or climatic drivers.

Interestingly, our findings suggest that the force of infection, but not the recent incidence of dengue, is correlated to Zika/microcephaly incidence. This finding underscores the extent to which, for immunizing diseases in endemic circulation, recent incidence may be poor metric of transmission. Immunity of the population in high or very high transmission settings can blunt the incidence so that places where transmission intensity is actually lower may experience incidences than are equal or higher. Metrics such as the force of infection, that quantify the risk among the susceptible population, are therefore better metrics of the underlying transmission potential in a given setting and can be derived from data that is commonly collected by surveillance systems. Furthermore, since FOI estimates are derived from age-specific incidence data, they are more robust to differences in surveillance efficiency than simple case based measures.

Similarly, we found no correlation between chikungunya incidence during the 2014/2015 epidemic and Zika/microcephaly incidence. Such correlation would be expected since chikungunya was introduced into the American continent in 2014 and the population was known to be fully susceptible prior to that date.^10,11^ Variability in case detection between different states/departments in Brazil and Colombia might be driving this finding. Alternatively, this lack of correlation could suggest more fundamental differences in the transmissibility of these viruses, such as differential competence of specific vector species^21^.

While we cannot rule out whether surveillance biases driven by resources or general concern of arboviruses contributed to the observed correlation between dengue and Zika, it is reassuring that such correlation was only evident for the force of infection dengue, but not for recent incidence. Surveillance biases would probably favor the latter. Given that both Zika and dengue are flaviviruses, immunological interactions in human hosts or vectors could also be contributing or strengthening the shared spatial distribution of transmission intensity observed here. ^22,23^ However, there is no evidence that important interactions take place between these pathogens and such interaction is not necessary to explain our result.

Our projections of the incidence and number of expected microcephaly cases in Colombia should be interpreted with caution. While the number of microcephaly cases reported weekly in Brazil seems to be declining, the microcephaly epidemic is not over and therefore we could be underestimating the true incidence in some settings. More importantly, Brazil is in the process of investigating all reported microcephaly cases but so far only 1345/5280 (25%) cases have been investigated and only 508 have been confirmed.^20,24^ Thus, the overall false positivity rate seems to be high (~62%) and the true incidences of microcephaly are likely to be lower than those reported. We present both unadjusted and adjusted projections, but expect the latter to change as the proportion of cases investigated increases over the coming weeks.

Findings from this study suggest that knowledge on the transmission intensity of dengue in different settings can be used to guide surveillance, testing and control efforts, and to anticipate the potential impact of the Zika epidemic in different populations. Even though the causal association between Zika and microcephaly has yet to be established, the observed correlation also provides a unique opportunity to predict the expected number of microcephaly cases in dengue endemic areas with autochthonous Zika transmission. This could be of significant utility right now, since our lack of understanding of the absolute risk of microcephaly among pregnancies affected by Zika infections limits our capacity to produce other types of projections. Surveillance and control efforts for Zika should prioritize areas where dengue transmission potential is known to be highest.

## References

1 Fauci AS, Morens DM. Zika Virus in the Americas--Yet Another Arbovirus Threat. N Engl J Med 2016; 374: 601–4.

2 Editorial. The next steps on Zika. Nature 2016; 530: 5–5.

3 World Health Organization (WHO). WHO statement on the first meeting of the International Health Regulations (2005)(IHR 2005) Emergency Committee on Zika virus and observed increase in neurological disorders and neonatal. 2016; published online Feb 1. These results also provided insight into the distinct mechanisms that might be driving (accessed Feb 22, 2016).

4 Schuler-Faccini L, Ribeiro EM, Feitosa IML, et al. Possible Association Between Zika Virus Infection and Microcephaly - Brazil, 2015. MMWR Morb Mortal Wkly Rep 2016; 65: 59–62.

5 Mlakar J, Korva M, Tul N, et al. Zika Virus Associated with Microcephaly. N Engl J Med 2016;: 160210140106006.

6 Instituto Nacional de Salud. Boletín epidemiológico semanal. 2016 http://www.ins.gov.co/boletin-epidemiologico/Boletn%20Epidemiolgico/2016%20Boletin%20epidemiologico%20semana%204.pdf.

7 Guzman MG, Kouri G. Dengue and dengue hemorrhagic fever in the Americas: lessons and challenges. J Clin Virol 2003; 27: 1–13.

8 Simmons CP, Farrar JJ, van Vinh Chau N, Wills B. Dengue. N Engl J Med 2012; 366: 1423–32.

9 Bhatt S, Gething PW, Brady OJ, et al. The global distribution and burden of dengue. Nature 2013; 496: 504–7.

10 Weaver SC. Arrival of chikungunya virus in the new world: prospects for spread and impact on public health. PLoS Negl Trop Dis 2014; 8: e2921.

11 Leparc-Goffart I, Nougairede A, Cassadou S, Prat C, de Lamballerie X. Chikungunya in the Americas. The Lancet 2014; 383: 514.

12 Centro de operacoes de emergencias em saude publica sobre microcefalias. Informe Epidemiológico N° 11 – Semana Epidemiológica (Se) 04/2016 (24 a 30/01/2016). 2016 http://portalsaude.saude.gov.br/images/pdf/2016/fevereiro/03/COES-Microcefalias–-Informe-Epidemiol–gico-11–-SE-04-2016–-02FEV2016–-18h51-VDP.pdf.

13 Ministerio da Saude. Sistema de informações hospitalares/SIH/SUS. http://tabnet.datasus.gov.br/cgi/deftohtm.exe?sih/cnv/mruf.def.

14 Instituto Nacional de Salud. Boletín Epidemiológico semanal. 2014 http://www.ins.gov.co/boletin-epidemiologico/Boletn%20Epidemiolgico/2014%20Boletin%20epidemiologico%20semana%2053.pdf.

15 Instituto Nacional de Salud. Vigilancia Rutinaria. http://www.ins.gov.co/lineas-de-accion/Subdireccion-Vigilancia/sivigila/Paginas/vigilancia-rutinaria.aspx (accessed Jan 5, 2016).

16 Cummings DAT, Iamsirithaworn S, Lessler JT, et al. The Impact of the Demographic Transition on Dengue in Thailand: Insights from a Statistical Analysis and Mathematical Modeling. PLoS Med 2009; 6: e1000139.

17 Grenfell BT, Anderson RM. The estimation of age-related rates of infection from case notifications and serological data. J Hyg (Lond) 1985; 95: 419–36.

18 R Foundation for Statistical Computing. R:anguage and environment for statistical computing. http://www.R-project.org/.

19 Imai N, Dorigatti I, Cauchemez S, Ferguson NM. Estimating Dengue Transmission Intensity from Sero-Prevalence Surveys in Multiple Countries. PLoS Negl Trop Dis 2015; 9: e0003719.

20 Centro de operacoes de emergencias em saude publica sobre microcefalias. Informe Epidemiológico N° 13 – Semana Epidemiológica (Se) 06/2016 (07 a 13/02/2016) Monitoramento Dos Casos De Microcefalia No Brasil. 2016 http://portalsaude.saude.gov.br/images/pdf/2016/fevereiro/17/coes-microcefalia-inf-epi-13-se06-2016.pdf.

21 Brady OJ, Golding N, Pigott DM, Kraemer MU. Global temperature constraints on Aedes aegypti and Ae. albopictus persistence and competence for dengue virus transmission. Parasit … 2014.

22 Althouse BM, Hanley KA. The tortoise or the hare? Impacts of within-host dynamics on transmission success of arthropod-borne viruses. Phil Trans R Soc Lond B 2015; 370: 20140299.

23 Calisher CH, Karabatsos N, Dalrymple JM, et al. Antigenic relationships between flaviviruses as determined by cross-neutralization tests with polyclonal antisera. J Gen Virol 1989; 70 ( Pt 1): 37–43.

24 Victora CG, Schuler-Faccini L, Matijasevich A, Ribeiro E, Pessoa A, Barros FC. Microcephaly in Brazil: how to interpret reported numbers? Lancet 2016; 387: 621–4.

